# Systematic Comparison of Indicator and Pathogenic Viruses Using High-Throughput qPCR Identifies Pepper Mild Mottle Virus as a Robust Indicator of Virus Removal in Wastewater Treatment

**DOI:** 10.64898/2026.05.30.728929

**Authors:** Shotaro Torii, Bikash Malla, Hiroki Ando, Masaaki Kitajima, Eiji Haramoto

**Affiliations:** Department of Urban Engineering, School of Engineering, The University of Tokyo, 7-3-1 Hongo, Bunkyo-ku, Tokyo 113-8656, Japan; Interdisciplinary Center for River Basin Environment, University of Yamanashi, 4-3-11 Takeda, Kofu, Yamanashi 400-8511, Japan; Division of Environmental Engineering, Faculty of Engineering, Hokkaido University, North 13 West 8, Kita-ku, Sapporo, Hokkaido 060-8628, Japan; Mel and Enid Zuckerman College of Public Health, University of Arizona, Tucson, Arizona 85724, United States; Research Center for Water Environment Technology, School of Engineering, The University of Tokyo, 7-3-1 Hongo, Bunkyo-ku, Tokyo 113-8656, Japan; The University of Tokyo Pandemic Preparedness, Infection and Advanced Research Center (UTOPIA), The University of Tokyo, 4-6-1 Shirokanedai, Minato-ku, Tokyo 108-0071, Japan

**Author notes:** Corresponding author: Shotaro Torii.

**Keywords:** High-Throughput Quantitative PCR, Tobamovirus, Virus, Wastewater Treatment

## Abstract

The selection of appropriate viral indicators for evaluating wastewater treatment performance remains challenging because candidate markers have rarely been compared systematically within a unified analytical framework. Here, we collected influent and effluent samples monthly for one year from two wastewater treatment plants in Japan and conducted, to our knowledge, the first comprehensive comparison of 19 viral targets and one protozoan target using high-throughput quantitative PCR. Pepper mild mottle virus (PMMoV) was consistently detected at high concentrations, showed limited seasonal variability, and exhibited an approximately 1.0 log_10_ reduction, comparable to those observed for pathogenic viruses. In contrast, *Carjivirus*, formerly known as crAssphage, was present at the highest concentrations but showed significantly greater reduction than pathogenic viruses. Tomato brown rugose fruit virus (ToBRFV), despite its high abundance and emerging recognition as a potential marker, exhibited pronounced seasonal fluctuations. Other *Tobamovirus* species, such as cucumber green mottle mosaic virus and tobacco mild green mosaic virus, exhibited similar removal but lower prevalence compared with PMMoV. Overall, PMMoV demonstrated the most balanced performance in terms of abundance, stability, and removal behavior, supporting its use as a robust indicator for monitoring virus removal in wastewater treatment.

## 1. Introduction

Enteric viruses are one of the major microbial contaminants that cause waterborne diseases. Adenovirus (AdV), caliciviruses including norovirus (NoV) and sapovirus (SaV), and enterovirus (EV) have been found to cause waterborne transmission and are key targets for managing the safety of drinking water (World Health Organization, 2022). These viruses enter water bodies through untreated sewage as well as the effluent of the typical wastewater treatment processes, such as the activated sludge process. It is thus important to monitor the level of viral abundance in wastewater and the removal efficiency of a wastewater treatment plant (WWTP) (Hata et al., 2013).

Conventional fecal indicator bacteria (FIB) are inadequate proxies for viral pollution and its reduction, as viruses are removed less efficiently during treatment and persist longer in aquatic environments (Boehm et al., 2018). This discrepancy has driven extensive efforts to identify viral indicators capable of replacing FIB for routine monitoring (Farkas et al., 2020).

Indicator viruses must satisfy the following criteria: they should (i) be more abundant than pathogenic viruses, (ii) exhibit minimal seasonal variability, and (iii) undergo similar or lesser reduction compared with pathogenic viruses during treatment processes (Pepper et al., 2014). Pepper mild mottle virus (PMMoV), a non-enveloped, single-stranded RNA plant virus belonging to the genus *Tobamovirus*, has emerged as one of the most extensively studied indicator candidates over the past decade (Kitajima et al., 2018; Miura et al., 2024). PMMoV has been found to be most abundant in human feces (Zhang et al., 2006) and is present at high concentrations in influent (up to 10^10^ genome copies per liter (gc/L)) and secondary effluent (up to 10^9^ gc/L) in WWTPs. Despite its rod-shaped morphology, distinct from the enteric viruses, PMMoV is removed to a comparable extent as AdV, NoV, and EV by the activated sludge process (Kitajima et al., 2014; Tandukar et al., 2020). However, the shedding load of PMMoV in feces was reported to differ greatly across individuals (Arts et al., 2023), necessitating a further confirmation of its limited variability in wastewater.

Advances in metagenomics have identified additional indicator viruses. One example is *Carjivirus* (formerly referred to as cross-assembly phage (crAssphage)), which belongs to the order *Crassvirales* and was discovered as double-stranded DNA bacteriophages highly abundant in the human gut (Dutilh et al., 2014; Yutin et al., 2018). A genetic marker of *Carjivirus*, CPQ_056 (Stachler et al., 2017), was found to be present at higher concentrations than enteric viruses in wastewater (Tandukar et al., 2020; Wu et al., 2020). Relative removal of *Carjivirus* to pathogenic viruses remains elusive: Farkas et al., 2019 reported similar reduction, while Tandukar et al. (2020) reported higher removal of *Carjivirus* compared with enteric viruses.

Other proposed indicators include *Tobamovirus* species (D’Aoust et al., 2026). For instance, a newly emerging plant virus, tomato brown rugose fruit virus (ToBRFV), was reported to be more abundant than PMMoV in feces and wastewater influent solids in the US (Natarajan et al., 2023; Sherchan et al., 2023) and was also detected in different countries such as Thailand (Paisantham et al., 2025). Cucumber green mottle mosaic virus (CGMMV) was found to be more abundant than PMMoV in WWTP influent in Slovenia (Bačnik et al., 2020) and in WWTP effluent in Arizona, the US (Yasui et al., 2021). Metagenomic studies reported high abundances of tobacco mild green mosaic virus (TMGMV), tomato mosaic virus (ToMV), tomato mottle mosaic virus (ToMMV), tropical soda apple mosaic virus (TSAMV), and tobacco mosaic virus (TMV) in wastewater and sewage-impacted environmental waters (Bačnik et al., 2020; Brumfield et al., 2022; Cuevas-Ferrando et al., 2025; Lopez-Roblero et al., 2023; Rosario et al., 2009; Rothman et al., 2023). However, to our knowledge, no studies have comprehensively compared PMMoV with other proposed indicators due to labor-intensiveness of repeating runs of single-plex quantitative PCR. It still remains unclear whether these Tobamoviruses fulfill the criteria of viral indicators and outperform PMMoV.

The high-throughput quantitative PCR (HT-qPCR) system offers a powerful tool to overcome this challenge. This system carries out thousands of real-time PCR reactions with nanoliter-scale per chamber in a single run by employing a nanofluid technology. In fact, HT-qPCR has been used to simultaneously quantify microbial source tracking markers and human-infectious pathogens in environmental water and wastewater (Hill et al., 2023; Ishii et al., 2014, 2013; Malla et al., 2022; Shrestha et al., 2024).

Here, we employed an HT-qPCR system and comprehensively compared the abundance and removal of pathogenic viruses, *Carjivirus*, PMMoV, and other Tobamoviruses at two WWTPs. Additionally, to further discuss the usefulness of the indicators, we included previously proposed DNA viral indicators, BK polyomavirus (BKPyV) and JC polyomavirus (JCPyV) (Bofill-Mas et al., 2000; Rachmadi et al., 2016), two respiratory viruses, including severe acute respiratory syndrome coronavirus 2 (SARS-CoV-2) and influenza A virus (IAV), and one protozoan, *Giardia* spp. as targets for the HT-qPCR. Finally, we proposed the best viral indicator based on its abundance, seasonal variability, and removal during the wastewater treatment process.

## 2. Materials and Methods

### 2.1. Sampling of influent and effluent of wastewater treatment plant

This study utilized the archived membrane samples obtained by filtering wastewater samples and storing the membranes in a freezer initially intended for wastewater-based epidemiological surveillance as a biobank (Kitajima et al., 2025). The samples of the influent and effluent from two WWTPs (Plants A and B) located in Sapporo, Japan, were collected monthly from February 2022 to January 2023. Both WWTPs employ the conventional activated sludge process, followed by chlorination. A total of 48 grab samples were collected, including 12 influent samples (after screening and before primary sedimentation) and 12 effluent (after chlorination and dechlorination) samples from each plant. All samples were collected in sterile plastic bottles, stored on ice, transported to the laboratory, and processed within 24 h of collection.

### 2.2. Concentration

Three hundred milliliters of influent or 2 L of effluent were supplemented with a final concentration of 25 mM MgCl_2_ and then filtered through an electronegative mixed cellulose-ester membrane (AAWP09000; pore size, 0.8 μm; diameter, 90 mm; Merck Millipore, Burlington, MA, USA). The filtered membranes were stored at -20°C at Hokkaido University and transported to the University of Tokyo prior to use.

### 2.3. Nucleic acid extraction and reverse transcription

DNA and RNA were co-extracted with a slightly modified version of the Efficient and Practical Virus Identification System with Enhanced Sensitivity for Membrane (EPISENS-M) (Ando et al., 2023b; Nakaso et al., 2024). This method was found to simultaneously concentrate and lyse protozoa, including *Cryptosporidium* and *Giardia*, and viruses with high efficiency (Torii et al., 2026). A quarter of the membrane was inserted into Precellys 7 mL tubes (Bertin Technologies, Montigny-le-Bretonneux, France), where beads from the RNeasy PowerWater Kit (Qiagen, Hilden, Germany), 850 μL of preheated PM1, 150 μL of TRIzol (Life Technologies, Carlsbad, CA, USA), and 10 μL of β-mercaptoethanol (Wako, Osaka, Japan) were added. Additionally, 2 μL of murine norovirus (MNV) was included as a molecular process control (MPC) (Haramoto et al., 2018). The tubes were then subjected to bead-beating three times at 10,000 rpm for 20 sec with 15 sec intervals using the Precellys Evolution tissue homogenizer (Bertin Technologies). After centrifugation at 10,000 × *g* for 3 min, 450 μL of supernatant was transferred to a rotor adapter using a QIAcube Connect platform (Qiagen). Co-extraction of DNA and RNA was performed with the RNeasy PowerMicrobiome Kit (Qiagen) according to the manufacturer’s instructions; inhibitor removal steps were included, but DNase treatment was not performed. DNA/RNA were extracted with a final volume of 50 μL of DNase/RNase free water.

Subsequently, 7.5 μL of the extract was subjected to reverse transcription (RT) using the High-Capacity cDNA Reverse Transcription Kit (Thermo Fisher Scientific, Waltham, MA, USA) to obtain 15 μL of (c)DNA, following the manufacturer’s protocol. The (c)DNA mixture was stored at -20°C and transported to the University of Yamanashi for further processing.

### 2.4. Pre-amplification

Pre-amplification, also known as specific target amplification, was carried out as described elsewhere (Ishii et al., 2013; Malla et al., 2022). Forward and reverse primers (20 μM each) for each assay were mixed in a single tube at a final concentration of 0.2 μM to prepare a pooled assay. Pre-amplification reactions (5 μL) were prepared by mixing 1.0 μL of PreAmp Master Mix (Standard BioTools, South San Francisco, CA, USA), 1.25 μL of the pooled assay, 1.5 μL of PCR-grade water, and 1.25 μL of DNA/cDNA, positive controls or negative controls, according to a previous study (Shrestha et al., 2024). As positive controls, ten-fold serial dilutions (1.0 × 10^4^ to 1.0× 10^-1^ genome copies (gc)/μL for SARS-CoV-2 and AdV group F (AdV-F) assays and 1.0 × 10^5^ to 1.0 × 10^0^ gc/μL for the others) of gBlocks or synthesized plasmid DNA were used. PCR-grade water was used as a negative control for pre-amplification. Pre-amplification was performed in a TaKaRa PCR Thermal Cycler Dice Touch (Takara Bio, Kusatsu, Japan) using the following thermal conditions: an initial activation at 95 °C for 2 min, followed by 14 cycles of denaturation at 95 °C for 15 s and annealing/extension at 60 °C for 4 min. The amplified products were diluted with TE buffer (cat. no. 12090015; Thermo Fisher Scientific) in a 1:5 ratio and immediately subjected to HT-qPCR.

### 2.5. High-throughput PCR (HT-qPCR)

A total of 23 different qPCR assays were initially tested to develop an HT-qPCR assay. These assays included two types for protozoa (*Cryptosporidium* spp. and *Giardia* spp.), seven for pathogenic viruses (AdV-F, Aichi virus (AiV), EV, SaV, NoV genogroup I (GI), NoV GII, and rotavirus (RV)), two for respiratory viruses (SARS-CoV-2 and IAV), eight for *Tobamovirus* species (CGMMV, PMMoV, TMV, TMGMV, ToMMV, ToMV, ToBRFV, and TSAMV), three for indicator DNA viruses (BKPyV, JCPyV, and *Carjivirus*), and one assay for MNV. All the assays, except for TSAMV, IAV, and RV, were performed according to previous studies (Balique et al., 2013; Boben et al., 2007; Dovas et al., 2010; Haramoto et al., 2013; Hongyun et al., 2008; Iturriza-Gómara et al., 2006; Jothikumar et al., 2021, 2009; Kageyama et al., 2003; Katayama et al., 2002; Kitajima et al., 2013, 2010; Ko et al., 2005; Lu et al., 2020; Maachi et al., 2022; Natarajan et al., 2023; Oka et al., 2006; Pal et al., 2006; Schoen et al., 2023; Shieh et al., 1995; Stachler et al., 2017; Yamaji et al., 2010; Zhang et al., 2006). All the assays used in the present study were TaqMan-based qPCR, and the sequences of forward primers, reverse primers, and probes for the other assays are listed in Table S1 in the Supplemental Information (SI). The assays for IAV and RV were slightly modified from Dovas et al., 2010 and Jothikumar et al., 2009, respectively, to cover possible sequence variations. For TSAMV, a forward primer TSAMV-F (5’–ATCCCTGCAGAYGAATTTGG–3’, corresponding to nt 688–707 in Okeechobee isolate [Accession No: KU659022]), a reverse primer TSAMV-R (5’–AATGGAAAGCGGCATAACAA–3’, corresponding to nt 757–738), and a ZEN probe TSAMV-P (5’-FAM-ACCTCGTCT/ZEN/CAGAAGTGCCGCC-IBFQ-3’, corresponding to nt 729 to 708) were developed in-house as detailed in supplemental text S1 of the SI.

HT-qPCR was performed on the Biomark^TM^ X9 System with 96.96 Dynamic Arrays^TM^ IFC (Standard BioTools) according to a previous study (Shrestha et al., 2024). The system can analyze up to 9,216 reaction chambers with 6.7 nL per chamber. In this study, 48 samples along with positive controls, negative controls, and MPC were measured for the 23 different qPCR assays in quadruplicate in a single run. The 10× assay mix and reaction mix were individually prepared as follows. Each 10× assay mix (30 μL) contained 15 μL of 2× Assay Loading Reagent (Standard BioTools) and primers and probes at a final concentration of 8 μM for each primer and 1 μM for each probe, except for PMMoV, for which the final concentration of each primer and probe was 7.0 μM and 1.0 μM, respectively. Each reaction mix (7.0 μL) contained 3.5 μL of TaqMan Fast Advanced Master Mix (Thermo Fischer Scientific), 0.35 μL of 20× GE Sample Loading Reagent (Standard BioTools), and 3.15 μL of 5-fold diluted pre-amplified DNA/cDNA. A negative control was prepared for both mixes by substituting primers and probe or preamplified (c)DNA with PCR-grade water. Subsequently, 5.0 μL each of the 10× assay mix and the reaction mix were applied to the assay and the sample inlets of IFC in quadruplicate and in singlicate, respectively. Then, the control line fluid was injected into the 96.96 IFC. The loaded IFC was transferred to the X9^TM^ Biomark System. The thermal cycling protocol consisted of initial incubation at 50 °C for 2 min, followed by a step at 95 °C for 10 min, and then subjected to 40 cycles comprising 95 °C for 15 s and 60 °C for 60 s (Malla et al., 2022). No amplification signals were observed for the negative control of pre-amplification or HT-qPCR. The amplification curve was analyzed using Standard Bio Tools Real-Time PCR Analysis 1.0.2.

### 2.6. Data analysis

All data analyses were performed using R 4.3.0 (R Core Team, 2019). The descriptive summary statistics, such as geometric mean and standard deviation for virus and protozoa concentration datasets and the recovery of molecular process control with left-censored observations, where a portion of measurements is below quantification limits, were computed by a robust regression on order statistics (ROS) with the ‘cenros’ function in the R package {NADA}, assuming a lognormal distribution to impute the left-censored observations prior to the computation of summary statistics (Helsel, 2011; Tang et al., 2024). This method, also called the multiple imputation method, was shown to provide consistent estimation of the mean for real-world data on enterovirus concentration in water compared with other methods, such as the substitution of left-censored data with limit of quantification (LOQ) /√2 (Canales et al., 2018).

Log_10_ reduction of viruses and protozoa was determined by the log_10_ concentration of influent minus the log_10_ concentrations of effluent. This calculation results in four types of log_10_ reduction datasets depending on the existence of unquantified data in influent and effluent samples. First, (i) no censored data were observed for *Carjivirus*, PMMoV, BKPyV, CGMMV, TMGMV, ToBRFV, and ToMV. Second, (ii) datasets partly containing only left- or right-censored reduction values were obtained for AdV-F and JCPyV. Third, (iii) datasets including both left-censored (i.e., less than *x* log_10_ reduction) and right-censored (i.e., more than *y* log_10_ reduction) were observed for AiV, SaV, SARS-CoV-2, TMV, ToMMV, and *Giardia.* Finally, (iv) only censored data or no data on reduction were obtained for datasets on EV, NoV GI, NoV GII, IAV, and TSAMV, which were excluded from analysis on reduction. The descriptive summary statistics of datasets (i) and (ii) were reported using ROS. For dataset (iii), uncensored values cannot be imputed because the percentile positions of the censored data are not uniquely assigned. The descriptive summary statistics were thus calculated only from uncensored values. The statistical comparison of the virus or protozoa reduction was performed with ANOVA followed by *post hoc* Tukey’s honestly significant difference (HSD) test.

## 3. Results

### 3.1. Development of an HT-qPCR assay for simultaneous quantification of viruses and protozoa

To develop an HT-qPCR system capable of simultaneously quantifying viruses and protozoa, a total of 23 assays were examined for their amplification signals in positive controls. The standard curves for each assay are shown in Figure 1. All assays, except for RV, showed clear amplification signals and a linear relationship between quantification cycle (Ct) values and template DNA amount. The RV assay was excluded in the subsequent steps. For the remaining 22 assays, R^2^ values were >0.92 and the LOQ ranged from 10^0.1-2.1^ gc/reaction as detailed in Table S2. Note that quantitative results for *Cryptosporidium* spp. are not discussed in the current manuscript because the assay produced unspecific amplification in nucleic acid extracts from environmental waters (Torii et al., 2026).

**Figure 1.**
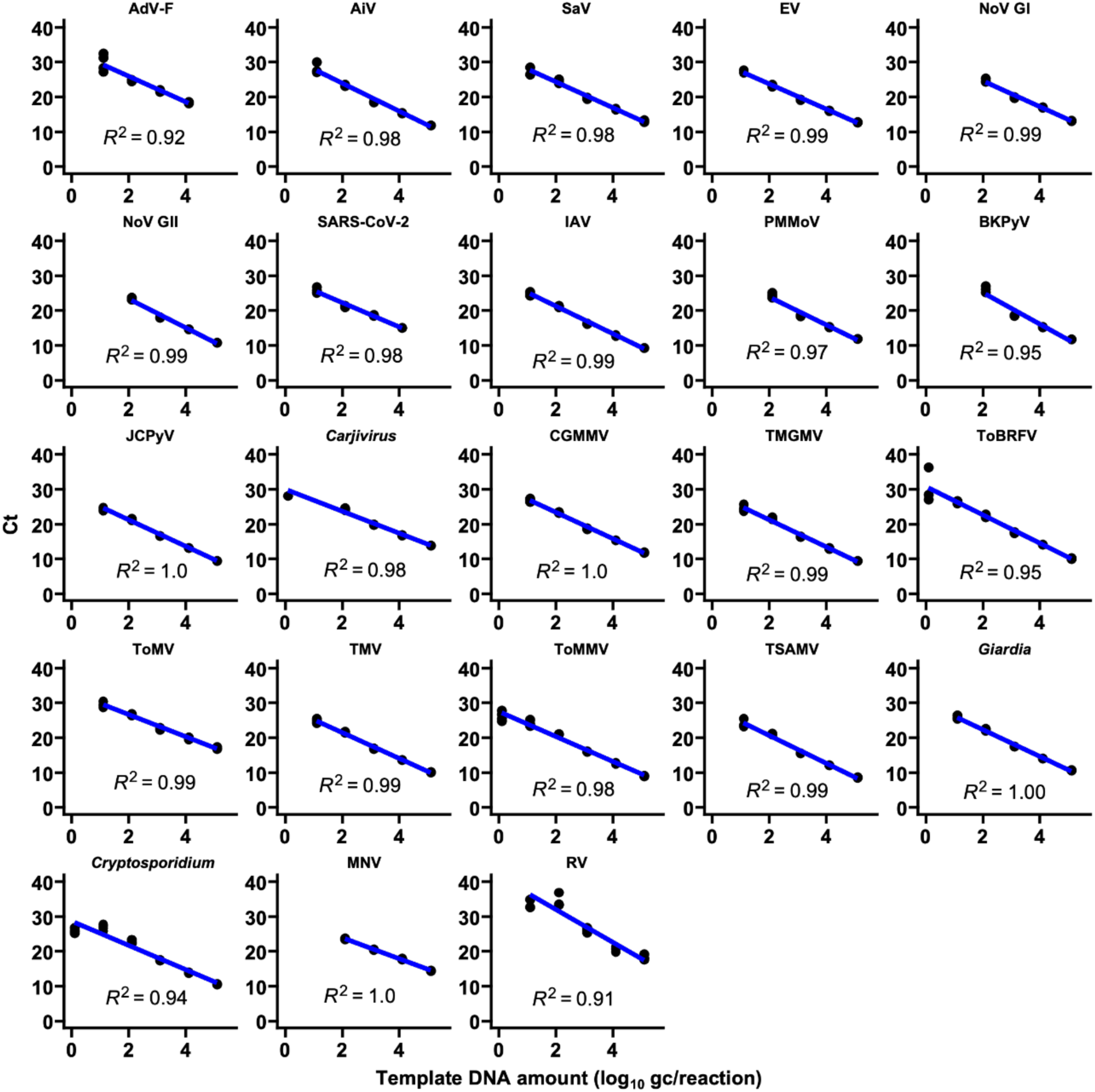
Standard curves for HT-qPCR assays used in the present study. Blue lines represent the standard curves generated by linear regression analysis of the quantification cycle (Ct) values obtained in HT-qPCR against the amounts of template DNA (log_10_ gc/reaction)

### 3.2. Recovery of molecular process control

To monitor the efficiency of molecular processes, including nucleic acid extraction, RT, pre-amplification, and HT-qPCR, MNV was spiked into the Precellys tubes. The MNV RNA concentration in the spiked control, which corresponded to 3.8 log_10_ gc/reaction, was determined based on the HT-qPCR quantification of the 100-fold diluted MNV stock extracted by the AllPrep PowerViral DNA/RNA Kit (Qiagen), using the same lysis and washing buffer used in the RNeasy PowerWater kit, followed by the same RT and preamplification process as the samples. The LOQ of MNV assay was determined to be 2.1 log_10_ gc/reaction, equivalent to 2.0% MPC recovery. MNV recovery was below the LOQ in 1/12 and 10/12 influent samples from Plants A and B, respectively. Using ROS, the mean MPC recovery (± SD) was estimated as follows: 56.1% ± 21.3% in Plant A influent, 29.6% ± 17.2% in Plant A effluent, 0.8% ± 1.5% in Plant B influent, and 79.0% ± 36.4% in Plant B effluent.

### 3.3. Abundance of viruses and protozoa in wastewater

A total of 48 samples were examined, consisting of 12 influent and 12 effluent samples from each of Plants A and B. These samples were analyzed by HT-qPCR for 19 viral targets, including human enteric viruses, respiratory viruses, viral indicators, and *Tobamovirus* species, as well as the protozoan target *Giardia*. The time-series concentrations are shown in Figure 2, and detailed summary statistics are provided in Table 1.

**Figure 2.**
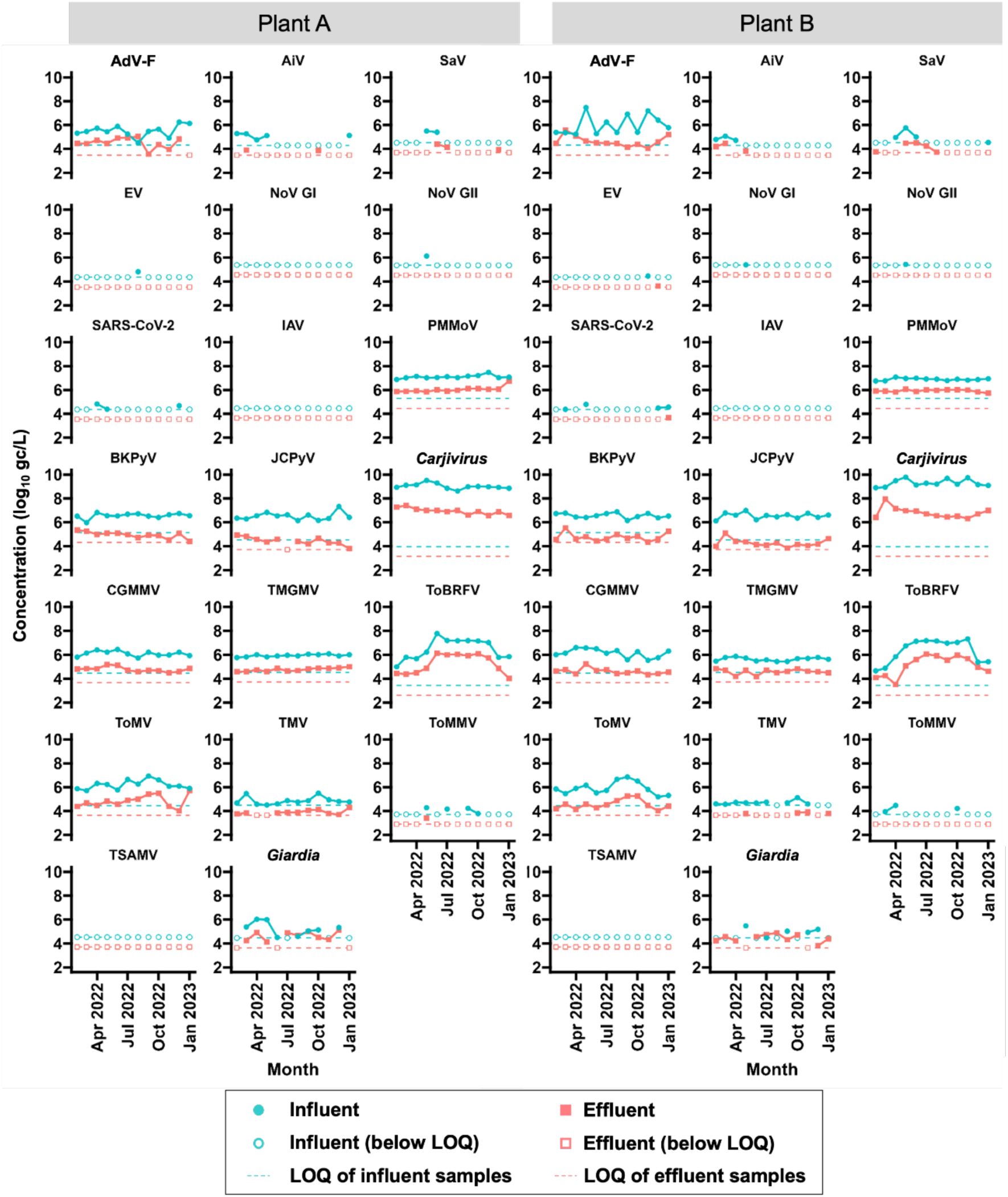
Time-series concentrations of enteric viruses, respiratory viruses, indicator viruses, *Carjivirus*, *Tobamovirus* species, and protozoa in influent and effluent samples (●, influent, ▪, effluent). The dashed lines represent the LOQ for both influent (blue) and effluent (red) samples.

**Table 1.**
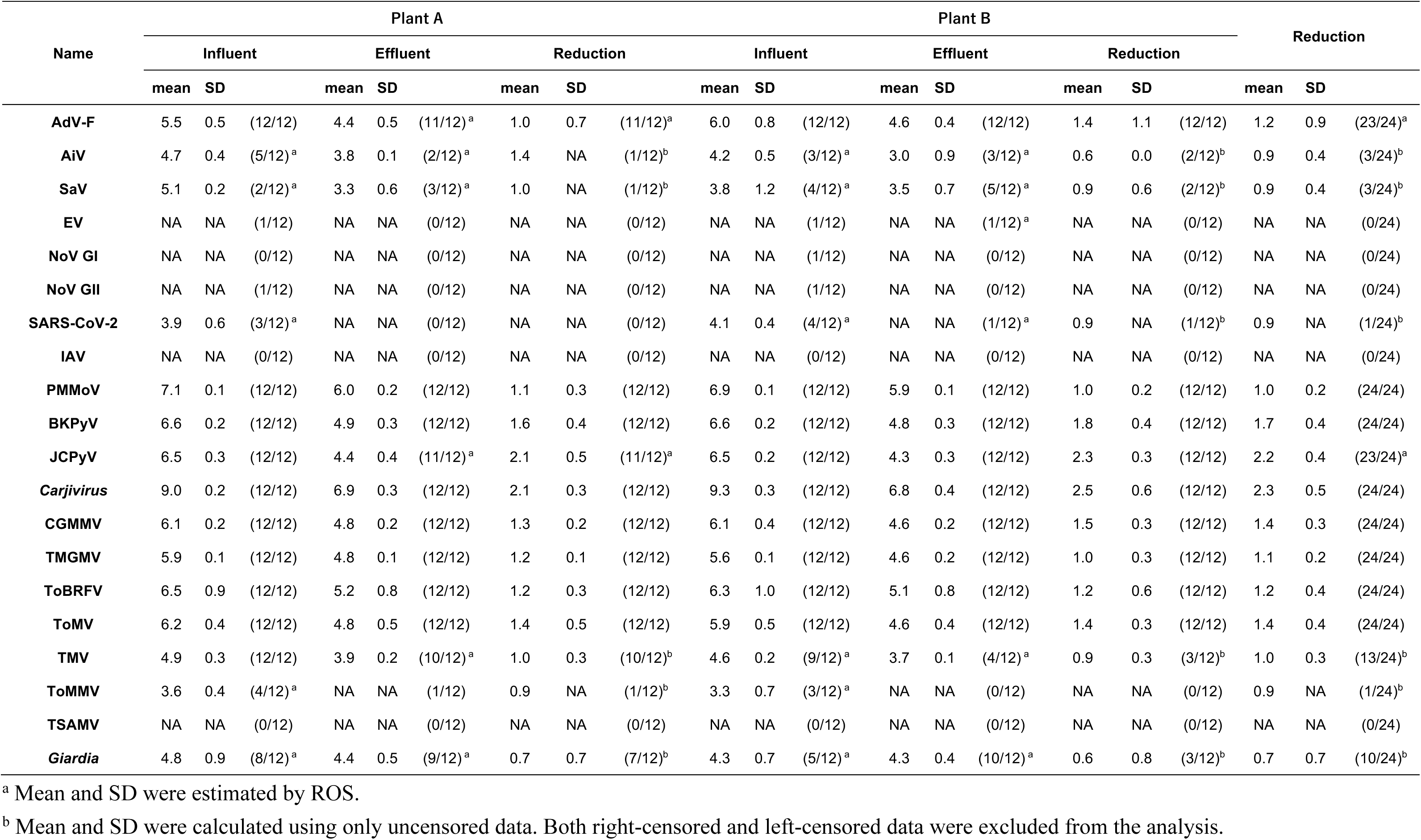
Arithmetic mean and standard deviation (SD) of log_10_ concentrations (log_10_ gc/L) and reduction of different viruses and protozoa. Reduction is expressed as the log-transformed ratio of effluent concentration to influent concentration (dimensionless). Numerators indicate the number of uncensored measurements while denominators represent the total number of samples in each plant.

Among human enteric viruses, AdV-F was consistently detected and showed the highest mean concentrations of 5.5–6.0 log_10_ gc/L in influent. In contrast, AiV and SaV were detected sporadically. EV, NoV GI, and NoV GII were rarely detected or not detected. Among the respiratory viruses, SARS-CoV-2 was detected in only a subset of influent samples but was absent from all effluent samples, whereas IAV was not detected in any samples.

Viral indicators, including *Carjivirus*, PMMoV, BKPyV, and JCPyV, were detected more consistently and at higher concentrations than the human enteric and respiratory viruses. *Carjivirus* showed the highest concentrations overall, reaching mean levels of 9.0–9.3 log_10_ gc/L in influent. PMMoV exhibited the second-highest mean concentration, 6.9–7.1 log_10_ gc/L.

Other *Tobamovirus* species were detected at concentrations lower than or comparable to PMMoV, but their detection patterns differed among targets. CGMMV and TMGMV were consistently detected at relatively high concentrations in both plants, whereas ToBRFV and ToMV showed clear seasonal patterns. In particular, ToBRFV increased from June to November 2022 and temporarily exceeded PMMoV concentrations during this period, but decreased substantially outside this season.

In addition to viruses, *Giardia* was detected in both influent and effluent samples from the two plants.

### 3.4. Reduction of viruses by wastewater treatment

To compare the removal of different viruses and protozoa, log_10_ reduction was calculated as the difference between influent and effluent concentrations. Data on EV, NoV GI, NoV GII, IAV, and TSAMV were excluded from the subsequent analyses because they either consisted of only censored data or no data on reduction. Time-series of viruses and protozoa reduction by each plant are shown in Figure S1. The descriptive summary statistics are shown in Table 1. None of the reductions in viruses and protozoa were found to be significantly different between the two plants (*P* > 0.05, *t*-test). Additionally, there was no observed seasonal trend in the reduction of viruses and protozoa. As a result, the reduction data from different plants and different time points were pooled and summarized into one dataset.

The comparison of the reduction of different viruses and protozoa is shown in Figure 3. The results of Tukey’s test are shown in Table S3. The reduction of PMMoV (1.0 ± 0.2 log_10_; n = 24) was similar to that of AdV-F (1.2 ± 0.9 log_10_; n = 23), AiV (0.9 ± 0.4 log_10_; n = 3), and SaV (0.9 ± 0.4 log_10_; n = 3). However, the other three indicator viruses, *Carjivirus*, BKPyV, and JCPyV, exhibited significantly higher reductions of 2.3 ± 0.5 log_10_ (n = 24), 1.7 ± 0.4 log_10_ (n = 24), and 2.2 ± 0.4 log_10_ (n = 24), respectively. Other *Tobamovirus* species, including CGMMV, TMGMV, ToBRFV, ToMV, and TMV, had similar reductions to those of PMMoV and enteric viruses, and the differences were not significant (*P* > 0.05, ANOVA followed by *post hoc* Tukey HSD’s test). The reduction in *Giardia* was 0.7 ± 0.7 log_10_, and were slightly lower than those observed for the viruses.

**Figure 3.**
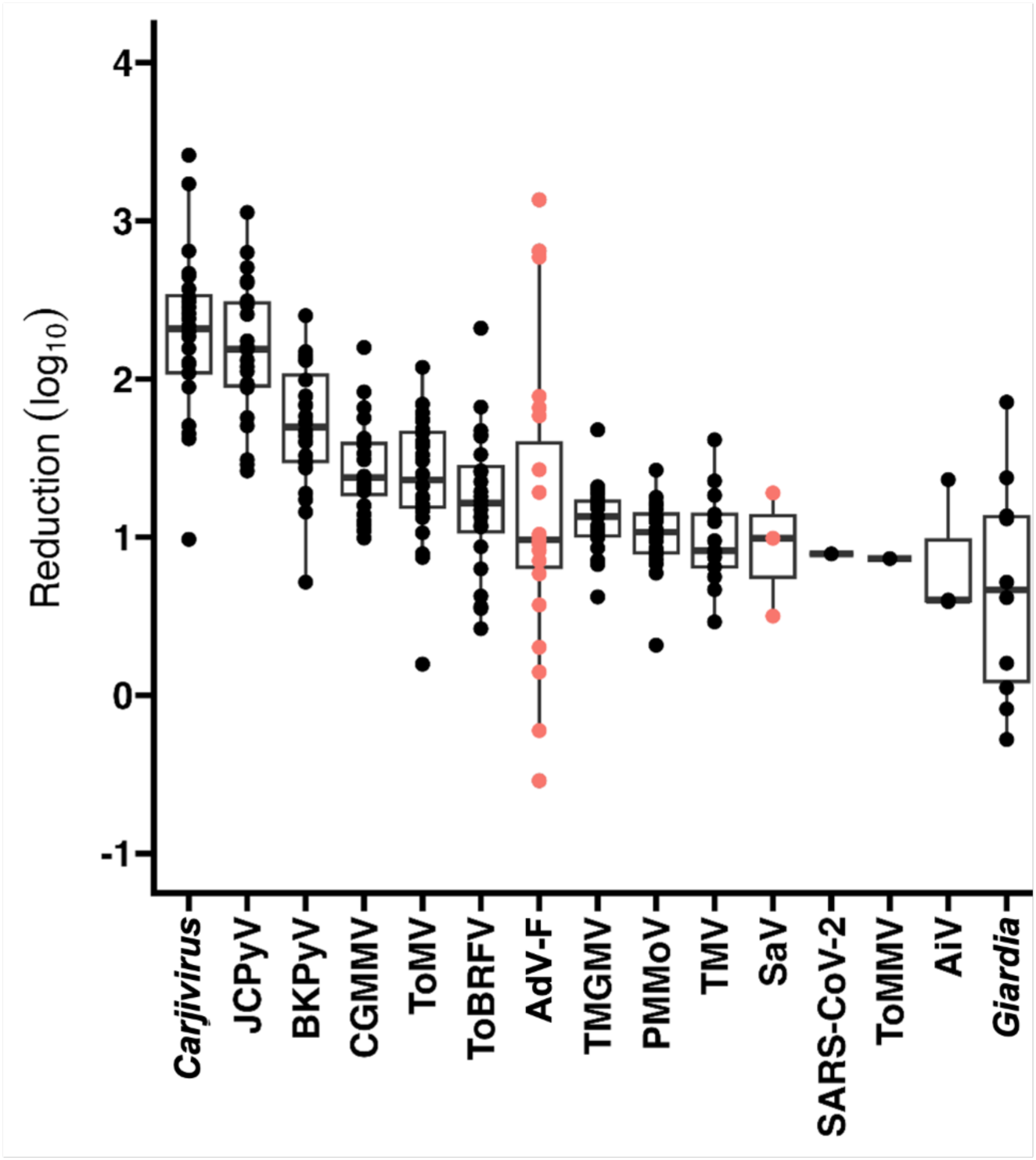
Comparative reduction of viruses and protozoa. Each plot represents different data points. Data on AdV-F and SaV were highlighted in red color. The lower whisker extends from the bottom of the box to the smallest value within 1.5 × interquartile range (IQR) of the lower quartile. The bottom edge of the box represents the first quartile. The line in the middle of the box represents the median. The top edge of the box represents the third quartile. The upper whisker extends from the top of the box to the largest value within 1.5 × IQR of the upper quartile.

### 3.5. Integrated evaluation of candidate viral indicators

To identify highly and stably prevalent viruses exhibiting similar removal to pathogenic viruses, the geometric mean was ranked in descending order, whereas coefficient of variation (CV) and the absolute differences of reduction values with AdV-F and SaV were ranked in ascending order (Figure 4 and Table S4).

**Figure 4.**
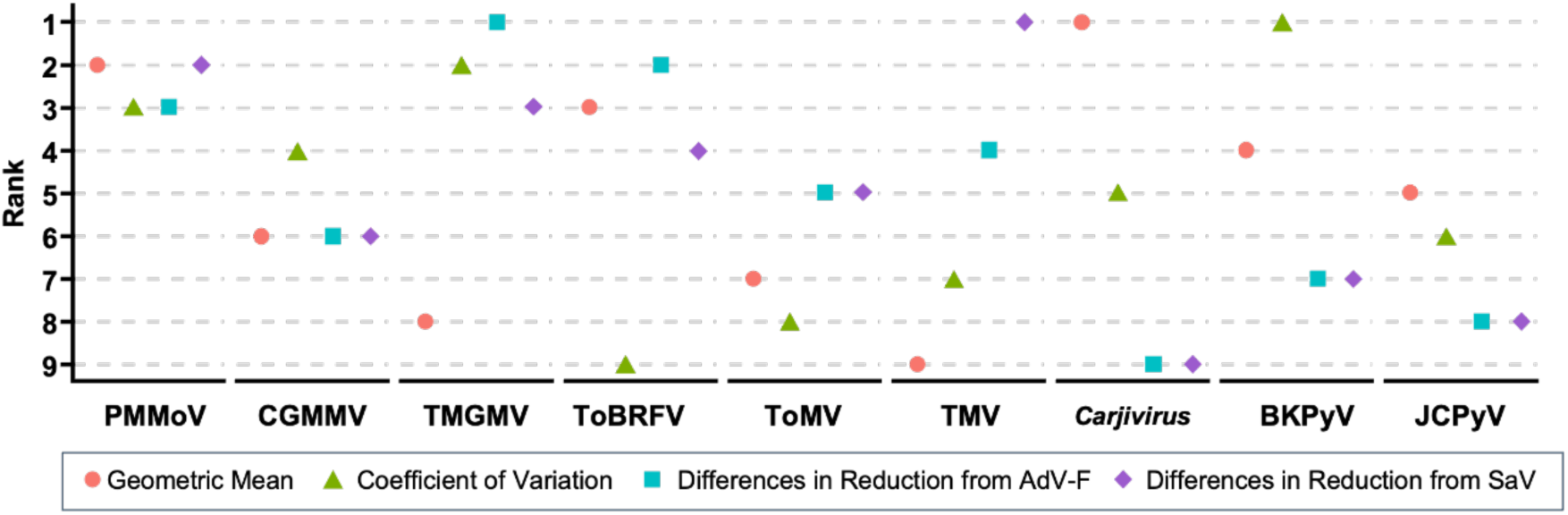
Descending rank of geometric mean, and the ascending rank of coefficient of variation and absolute differences in log_10_ reductions with AdV-F and SaV among nine viruses, including PMMoV CGMMV, TMGMV, ToBRFV, ToMV, TMV, *Carjivirus*, BKPyV, and JCPyV. Higher ranks (lower numbers) indicate more favorable characteristics as candidate viral indicators.

PMMoV showed the second-highest geometric mean concentration, the third lowest CV, as well as the third- and second-smallest differences in reduction from AdV-F and SaV, respectively, ranging from -0.1 log_10_ to 0.1 log_10_. ToBRFV exhibited the third-highest concentration and comparable removal to AdV-F and SaV; however, the variation across the season was the largest among the targets examined. Other types of *Tobamovirus* species, such as CGMMV, TMGMV, ToMV, and TMV, demonstrated comparable removal with AdV-F and SaV.

*Carjivirus* exhibited the highest geometric mean concentration, surpassing the second highest PMMoV by 2.1 log_10_. However, *Carjivirus* was removed at higher levels than AdV-F by 1.1 log_10_ and SaV by 1.4 log_10_. Similarly, BKPyV and, to a lesser extent, JCPyV were removed more than AdV-F and SaV.

## 4. Discussion

Using HT-qPCR, this study provided time-series data on the abundance and reduction of human pathogenic viruses, indicator viruses, *Tobamovirus* species, and protozoa at WWTPs in a unified analytical framework. The observed removal was based on genome copies and can be interpreted to be mostly attributed to the activated sludge process, as chlorination of secondary effluent in Japan was found to result in minimal decay of viral genomes measured with qPCR (Hata et al., 2013; Singhopon et al., 2026). In accordance with a previous study (Katayama et al., 2008), all types of enteric viruses except for AdV-F were detected during only certain periods throughout the year. This once again highlights the importance of the viral indicators.

The systematic comparisons of nine viral indicators identified PMMoV as a balanced marker for assessing enteric virus removal by wastewater treatment due to its high abundance, low seasonal variation, and comparable removal with pathogenic viruses.

These findings are consistent with previous studies reporting its superiority as an indicator (Kitajima et al., 2014; Tandukar et al., 2020).

*Carjivirus* was less suitable as an indicator because of its higher removal relative to pathogenic viruses (Tandukar et al., 2020; Wu et al., 2020). The higher removal of *Carjivirus* may be attributable to its unique biological lifestyle. A recent study revealed that the prototypic *Carjivirus* predominantly exists within its host as a linear phage-plasmid and rarely transitions into a lytic phase (Schmidtke et al., 2025), suggesting that the dominant portion of measured *Carjivirus* originates from the intracellular fraction within host cells. This is consistent with findings that *Carjivirus* is abundantly detected in the solid fraction of environmental waters, such as solids trapped by microfiltration membrane (Ballesté et al., 2019; Chen et al., 2019). These findings suggest that reductions in *Carjivirus* may primarily reflect removal of its host bacteria rather than removal of free phage-form *Carjivirus*, and thus may not be a suitable indicator for virus removal. 2026

Similarly, BKPyV and JCPyV were also highly prevalent in influent, consistent with previous studies, but exhibited higher removal, consistent with previous studies (Hata et al., 2013; Kitajima et al., 2014; Schmitz et al., 2016; Wu et al., 2020). The higher removal is also attributable to the physical-state of PyV. Both types of PyV are known to persist latently in the kidney and urinary tract (Boldorini et al., 2005; Singh et al., 2006) and can thus be excreted in a cell-associated state, implying that a certain proportion of PyVs are cell-associated in wastewater influent. Further understanding of solid-liquid partitioning would explain the differential removal of these viruses.

ToBRFV, despite attracting growing interest as an emerging indicator, was disqualified by its pronounced seasonal variation in concentration. This seasonality is partly consistent with reports from Louisiana, the US, where quantitative ToBRFV data in wastewater influent also showed peak concentrations around September to December (Sherchan et al., 2023). A contrasting study reported a constantly high concentration of ToBRFV in influent wastewater from different places in the US, although their samples were mostly collected in June and July of different years (Natarajan et al., 2023). The drivers of the seasonal difference need to be investigated in further research. ToBRFV was first reported in 2015 in Jordan and currently causes pandemic disease outbreaks among tomato and pepper crops worldwide. Notably, ToBRFV infection is not officially reported in Japan as of May 2026 (European and Mediterranean Plant Protection Organization (EPPO), 2026; Salem et al., 2023) despite its detection in wastewater in the present study. This discrepancy can be explained by possible contribution of imported vegetables or processed foods. Nevertheless, tracking the source of ToBRFV detected in Japanese wastewater remains an important area for future research.

Other *Tobamovirus* species, including CGMMV, TMGMV, ToMV, and TMV, showed removal values similar to PMMoV but were less abundant. Among them, ToMV also exhibited relatively large temporal variability. It should be noted that the calculated CV should be interpreted with caution because it reflects not only true inter-sample variation, but also random analytical error associated with HT-qPCR. In particular, random error is expected to increase when target concentrations are low; therefore, the CVs of targets with lower geometric mean concentrations, such as ToMV and TMV, may have been overestimated. Nevertheless, their lower abundance makes them less advantageous than PMMoV as indicators.

The present study has several limitations. First, additional validation is required to evaluate the efficiency of nucleic acid extraction from membranes. Although liquid MNV stock was externally inoculated onto part of the membrane as an MPC, this approach may not fully represent the lysis and recovery efficiency of viruses retained on the membrane. This study did not adjust the HT-qPCR measurements for recovery efficiency, which may have underestimated the absolute concentration of each target, particularly for Plant B influent samples, where MNV recovery was frequently below commonly used recovery thresholds of 1% (Haramoto et al., 2018). Further studies are needed to determine whether the lower recovery stems from insufficient lysis, PCR inhibition, or an artifact associated with MNV inoculation.

Second, although this study showed similarities in average removal levels, it could not sufficiently evaluate correlations between the reductions of candidate indicator viruses and enteric viruses because NoV GI, NoV GII, and EV were rarely quantified. This limited quantification may be attributable to the sensitivity constraints of the HT-qPCR workflow, given that a previous wastewater-based epidemiological surveillance study conducted at the same WWTPs during the same period, using RT-preamplification followed by single-plex RT-qPCR, detected SaV and AiV more frequently (Ando et al., 2023a). Possible countermeasures include increasing the template volume and optimizing the preamplification step. For targets that are already abundant enough to be quantified by nanoliter-scale HT-qPCR, omitting preamplification, as done for PMMoV by Ando et al., 2023a, may reduce competition for reaction components (e.g., dNTPs) and thereby facilitate the amplification of low-abundance targets. Such optimization would increase the number of quantifiable pathogenic-virus datasets and enable robust evaluation of whether the indicator viruses respond similarly to enteric viruses even when treatment performance varies.

Finally, the relative abundance and temporal stability of indicator viruses should be further evaluated across diverse geographic regions, where dietary habits, agricultural products, and imported foods may differ substantially. Such validation is needed to determine whether the superior performance of PMMoV observed in this study is broadly applicable.

## 5. Conclusion

The HT-qPCR-based systematic comparison revealed that PMMoV fulfilled the key criteria for a viral indicator in wastewater treatment: high prevalence, limited seasonal variation, and a removal comparable to that of pathogenic viruses. In contrast, recently proposed viral indicators, including *Carjivirus* and ToBRFV, have specific limitation; *Carjivirus* undergoes higher reduction during the treatment process while ToBRFV has a pronounced seasonal fluctuation despite its high prevalence. Overall, PMMoV is the most balanced surrogate for pathogenic virus removal among the candidate markers examined. Routine monitoring of PMMoV could therefore support more reliable evaluation of virus removal performance in activated sludge-based wastewater treatment.

## 6. Acknowledgments

This study was supported by AMED under Grant Number JP223fa627001 (UTOPIA Next-Generation Diagnostic Tool Discovery Program) and 26fk0108713h0003, the River Fund of The River Foundation, Japan under Grant Number 2023-5311-005, Kurita Water and Environmental Foundation, Next Generation Water Supply Research Incentive Program for Young Researchers supported by Kubota, the Japan Society for the Promotion of Science (JSPS) through Grant-in-Aid for Scientific Research (A) (grant number JP24H00326 and JP25H00755), Grant-in-Aid for Scientific Research (B) (grant number JP23H01536 and JP26K01072), Grant-in-Aid for Young Scientists (grant number JP24K17379), the Environment Research and Technology Development Fund (grant number JPMEERF20255RB1 and JPMEERF20265001) of the Environmental Restoration and Conservation Agency provided by Ministry of the Environment of Japan and Grant-in-Aid for JSPS Fellows (grant number JP23KF0036), the Japan Science and Technology Agency (JST) through e-ASIA Joint Research Program (grant number JPMJSC20E2). We acknowledge Standard BioTools K.K. for supporting our experiment using Biomark X9.

## 7. CRediT authorship contribution statement

**Shotaro Torii:** Formal analysis, Conceptualization, Funding acquisition, Investigation, Methodology, Validation, Writing – Original draft, **Bikash Malla**: Investigation, Methodology, Writing – review and editing, **Hiroki Ando:** Investigation, Methodology, Writing – review and editing, **Masaaki Kitajima:** Funding acquisition, Writing – review and editing, **Eiji Haramoto:** Funding acquisition, Writing – review and editing

## 8. Declaration of generative AI and AI-assisted technologies in the manuscript preparation process

During the preparation of this work the authors used ChatGPT solely for improving readability and language in accordance with the journal’s policies. After using this tool, the authors reviewed and edited the content as needed and take full responsibility for the content of the published article.

