## Supplementary material for "Systematic Comparison of Indicator and Pathogenic Viruses Using High-Throughput qPCR Identifies Pepper Mild Mottle Virus as a Robust Indicator of Virus Removal in Wastewater Treatment": SI

### Supplemental Text

#### S1: Development of RT-qPCR assay for the detection of tropical soda apple mosaic virus

The eleven nearly full-length nucleotide sequences of tropical soda apple mosaic virus (TSAMV) (Accession No. KU659022, MT130387, MT130386, MT130385, MT130383, MT130384, MT130379, MT130381, MT130382, MT130380, NC_030229) were aligned in Geneious Prime 2023.1.2 using the MAFFT plugin with default settings. The primers and probe sequences were designed based on highly conserved regions and identified using Real-time PCR (TaqMan) Primer and Probes Design Tool (GenScript; https://www.genscript.com/tools/real-time-pcr-taqman-primer-design-tool ).

The specificity of the assay was investigated *in silico*. First, a nucleotide BLAST search revealed that no sequences, except for TSAMV, completely matched the primers and probe sequences. To assess the homology with other *Tobamovirus* species, the sequences of 33 genetically-close species were aligned: Bell pepper mottle virus (DQ355023), Brugmansia mild mottle virus (AM398436), Cactus mild mottle virus (EU043335), Clitoria yellow mottle virus (JN566124), Cucumber fruit mottle mosaic virus (AF321057), Cucumber green mottle mosaic virus (D12505), Cucumber mottle virus (AB261167), Frangipani mosaic virus (HM026454), Hibiscus latent Fort Pierce virus (AB917427), Hibiscus latent Singapore virus (AF395898), Kyuri green mottle mosaic virus (AJ295948), Maracuja mosaic virus (DQ356949), Obuda pepper virus (D13438), Odontoglossum ringspot virus (X82130), Paprika mild mottle virus (AB089381), Passion fruit mosaic virus (HQ389540), Pepper mild mottle virus (M81413), Plumeria mosaic virus (KJ395757), Rattail cactus necrosis-associated virus (JF729471), Rehmannia mosaic virus (EF375551), Ribgrass mosaic virus (HQ667979), Streptocarpus flower break virus (AM040955), Tobacco mild green mosaic virus (M34077), Tobacco mosaic virus (V01408), Tomato brown rugose fruit virus (KT383474), Tomato mosaic virus (AF332868), Tomato mottle mosaic virus (KF477193), Tropical soda apple mosaic virus (KU659022), Turnip vein-clearing virus (U03387), Wasabi mottle virus (AB017503), Yellow tailflower mild mottle virus (KF495564), Youcai mosaic virus (U30944), and Zucchini green mottle mosaic virus (AJ295949). At least three, one, and five mismatches were observed in the forward primer, reverse primer, and TaqMan probe regions, respectively. Based on these findings, the developed assay is specific to TSAMV *in silico*.

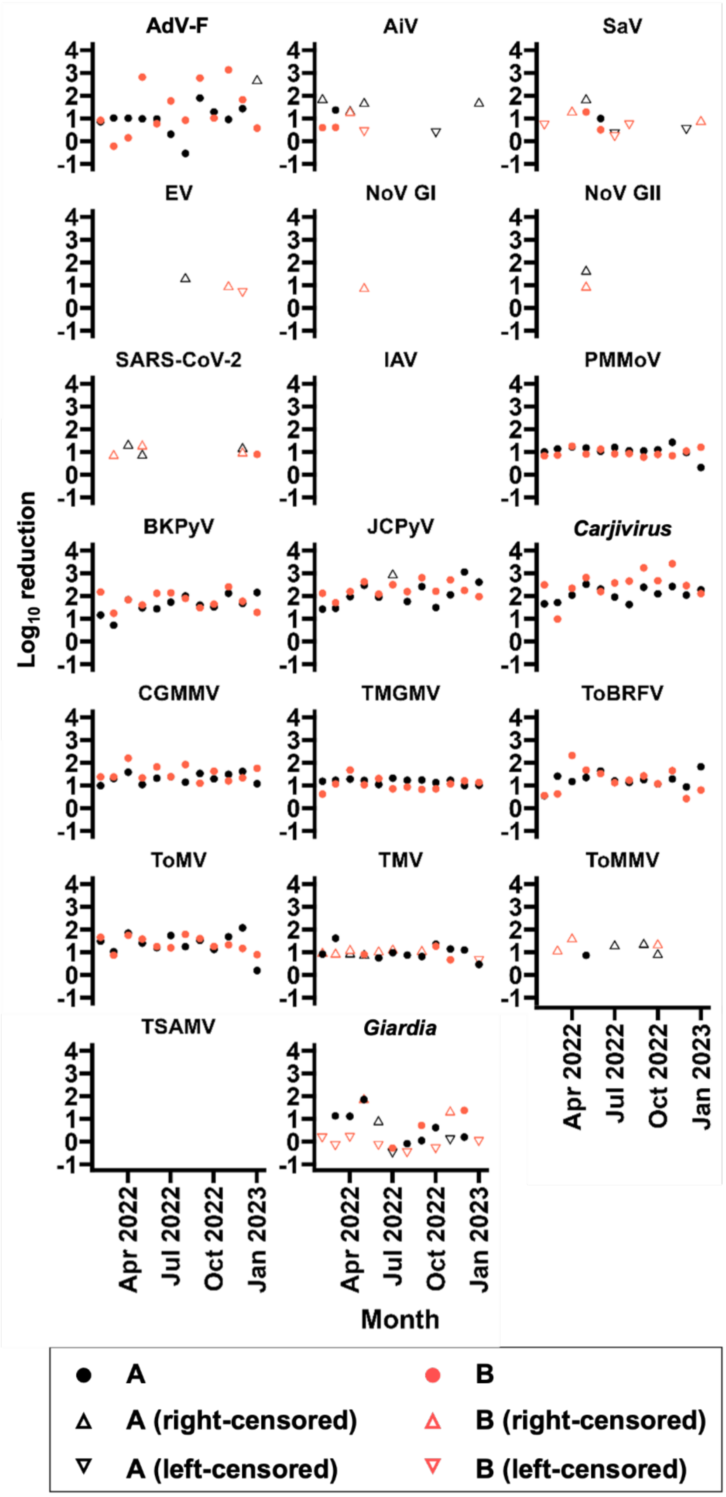


Figure S1. Time-series of virus and protozoa reduction by wastewater treatment. Uncensored, right-censored, and left-censored data were plotted with ●(closed circle), △ (open up-pointing triangle), and ▽ (open down-pointing triangle), respectively. Data for plant A and plant B are plotted with black and red color, respectively.

Table S1. Primers and probes for HT-qPCR.

| Target | Function | Name | Sequence (5' – 3')[a] | Amplicon length | Polarity | Reference |
|---|---|---|---|---|---|---|
| AdV-F | F | JTVFF | AACTTTCTCTCTTAATAGACGCC | 100 | + | (Ko et al., 2005) |
| AdV-F | R | JTVFR | AGGGGGCTAGAAAACAAAA | | - | |
| AdV-F | P | JTVFP | FAM-CTGACACGGGCACTCTTCGC-TAMRA | | + | |
| AiV | F | AiV-AB-F | GTCTCCACHGACACYAAYTGGAC | 108 | + | (Kitajima et al., 2013) |
| AiV | R | AiV-AB-R | GTTGTACATRGCAGCCCAGG | | - | |
| AiV | P | AiV-AB-TP | FAM-TTYTCCTTYGTGCGTGC-MGB-NFQ | | + | |
| SaV | F | SaV124F | GAYCASGCTCTCGCYACCTAC | 107 | + | (Oka et al., 2006) |
| SaV | F | SaV1F | TTGGCCCTCGCCACCTAC | | + | |
| SaV | F | SaV5F | TTTGAACAAGCTGTGGCATGCTAC | | + | |
| SaV | R | SaV1245R | CCCTCCATYTCAAACACTA | | - | |
| SaV | P | SaV124TP | FAM-CCRCCTATRAACCA-MGB-NFQ | | - | |
| SaV | P | SaV5TP | FAM–TGCCACCAATGTACCA-MGB-NFQ | | - | |
| EV | F | EntF (EV-L) | CCTCCGGCCCCTGAATG | 154 | + | (Shieh et al., 1995) |
| EV | R | EntR.v2 | ATTGTCACCATAAGCAGCCA | | - | (Iturriza-Gómara et al., 2006) |
| EV | P | Pan-Entero-P | FAM-CCGACTACTTTGGGTGTCCGTGTTTC-BHQ1 | | + | (Katayama et al., 2002) |
| NoV GI | F | COG1F | CGYTGGATGCGNTTYCATGA | 85 | + | (Kageyama et al., 2003) |
| NoV GI | R | COG1R | CTTAGACGCCATCATCATTYAC | | - | |
| NoV GI | P | RING1(a)-TP | FAM-AGATYGCGATCYCCTGTCCA-BHQ1 | | - | |
| NoV GI | P | RING1(b)-TP | FAM-AGATCGCGGTCTCCTGTCCA-BHQ1 | | - | |
| NoV GII | F | COG2F | CARGARBCNATGTTYAGRTGGATGAG | 98 | + | (Kageyama et al., 2003) |
| NoV GII | R | COG2R | TCGACGCCATCTTCATTCACA | | - | |
| NoV GII | P | RING2-TP | FAM-TGGGAGGGCGATCGCAATCT-BHQ1 | | + | |
| SARS-CoV-2 | F | 2019-nCoV_N1-F | GACCCCAAAATCAGCGAAAT | 72 | + | (Lu et al., 2020) |
| SARS-CoV-2 | R | 2019-nCoV_N1-R | TCTGGTTACTGCCAGTTGAATCTG | | - | |
| SARS-CoV-2 | P | 2019-nCoV_N1-P | ACCCCGCATTACGTTTGGTGGACC | | + | |
| IAV | F | InA/M-F | AGACCRATCYTGTCACCTCTGAC | 92 | + | (Dovas et al., 2010) |
| IAV | R | InA/M-R.v2 | AAKCGTCTACGCTGCAGTC | | - | This study |
| IAV | P | IAV-M_Neo_Probe | FAM-TTTGTGTTC/ZEN/ACGCTCACCGTGCCC-IBFQ | | + | This study |
| Carjivirus | F | 056F1 | CAGAAGTACAAACTCCTAAAAAACGTAGAG | 126 | + | (Stachler et al., 2017) |
| Carjivirus | R | 056R1 | GATGACCAATAAACAAGCCATTAGC | | - | |
| Carjivirus | P | 056P1 | FAM-AATAACGATTTACGTGATGTAAC-MGB-NFQ | | + | |
| PMMoV | F | PMMV-FP1-rev | GAGTGGTTTGACCTTAACGTTTGA | 68 | + | (Haramoto et al., 2013) |
| PMMoV | R | PMMV-RP1 | TTGTCGGTTGCAATGCAAGT | | - | (Zhang et al., 2006) |
| PMMoV | P | PMMV-Probe1 | FAM-CCTACCGAAGCAAATG-MGB-NFQ | | + | (Zhang et al., 2006) |

| | | | | | | |
|---|---|---|---|---|---|---|
| BKPyV | F | BKPyV-F | GGCTGAAGTATCTGAGACTTGGG | 78 | + | (Pal et al., 2006) |
| BKPyV | R | BKPyV-R | GAAACTGAAGACTCTGGACATGGA | | - | |
| BKPyV | P | BKPyV-TP | FAM-CAAGCACTG/ZEN/AATCCCAATCACAATGCTC-IBFQ | | - | |
| JCPyV | F | JCPyV-F | GGAAAGTCTTTAGGGTCTTCTACCTTT | 85 | + | (Pal et al., 2006) |
| JCPyV | R | JCPyV-R | ATGTTTGCCAGTGATGATGAAAA | | - | |
| JCPyV | P | JCPyV-TP | FAM-AGGATCCCA/ZEN/ACACTCTACCCCACCTAAAAAGA-IBFQ | | - | |
| CGMMV | F | CGMMV_F | GCATAGTGCTTTCCCGTTCAC | 101 | + | (Hongyun et al., 2008) |
| CGMMV | R | CGMMV_R | TGCAGAATTACTGCCCATAGAAAC | | - | |
| CGMMV | P | CGMMV_P | FAM-CGGTTTGCTCATTGGTTTGCGGA-BHQ1 | | + | |
| TMGMV | F | TMGMV-F | GTAGCTATAAGGGCTTCAATCA | 106 | + | (Maachi et al., 2022) |
| TMGMV | R | TMGMV-R | TGGTCCARACAAGTCCACTA | | - | |
| TMGMV | P | TMGMV-P | FAM-TGAACTGGT/ZEN/TCGTGGAACTGGCAT-IBFQ | | + | |
| ToBRFV | F | ToBRFV_Mo_F | TCAGTGTCTGTTTGGTCGATAA | 106 | + | (Natarajan et al., 2023) |
| ToBRFV | R | ToBRFV_Mo_R | GGAACGACTTTGAACTGAAACC | | - | |
| ToBRFV | P | ToBRFV_Mo_P | FAM-AGAGCGGAC/ZEN/GAGGCAACTCTTG-IBFQ | | + | |
| ToMV | F | ToMVF | TTGCCGTGGTGGTGTGAGT | 72 | + | (Boben et al., 2007) |
| ToMV | R | ToMVR | GACCCCAGTGTGGCTTCGT | | - | |
| ToMV | P | ToMVMGB | FAM-TCTTTCCATTCTCTTGTCAA-NFQ-MGB | | - | |
| TMV | F | TMV_Mars_CPFwd1.v2 | CAAGCTCGAACTGTCGTTCA | 120 | + | (Yamaji et al., 2010) |
| TMV | R | TMV_Mars_CPRev1 | CGGGTCTAAYACCGCATTGT | | - | (Balique et al., 2013) |
| TMV | P | TMV_Mars_CP1 | FAM-CAGTGAGGTGTGGAAACCTTCACCACA-BHQ1 | | + | (Balique et al., 2013) |
| ToMMV | F | ToMMV CaTa 9-F | ATGTGGAGGAACCCTCTATGA | 105 | + | (Schoen et al., 2023) |
| ToMMV | R | ToMMV CaTa 9-R | AATCTCCTCGCTCCTTGTAAAC | | - | |
| ToMMV | P | ToMMV CaTa 9-P | FAM-TCAATGGCC/ZEN/CGTGGTGAGTTACAA-IBFQ | | + | |
| TSAMV | F | TSAMV-Fw | ATCCCTGCAGAYGAATTTGG | 70 | + | This study |
| TSAMV | R | TSAMV-Rv | AATGGAAAGCGGCATAACAA | | - | This study |
| TSAMV | P | TSAMV-Probe | FAM-ACCTCGTCT/ZEN/CAGAAGTGCCGCC-IBFQ | | - | This study |
| *Giardia* | F | 18SJVGF | ATCCGGTCGATCCTGCCG | 156 | + | (Jothikumar et al., 2021) |

| | | | | | | |
|---|---|---|---|---|---|---|
| *Giardia* | R | 18SJVGR | ACGTCTTGGCGCCGGGTT | | - | |
| *Giardia* | P | 18SJVGP | FAM-CGGCGGACG/ZEN/GCTCAGGA-IBFQ | | + | |
| *Cryptosporidium* | F | JVAF | ATGACGGGTAACGGGGAAT | 139 | + | (Jothikumar et al., 2008) |
| *Cryptosporidium* | R | JVAR | CCAATTACAAAACCAAAAAGTCC | | - | |
| *Cryptosporidium* | P | JVAP18S | FAM-CGCGCCTGC/ZEN/TGCCTTCCTTAGATG-IBFQ | | + | |
| MNV | F | MNV-S | CCGCAGGAACGCTCAGCAG | 127 | + | (Kitajima et al., 2010) |
| MNV | R | MNV-AS | GGYTGAATGGGGACGGCCTG | | - | |
| MNV | P | MNV-TP | FAM-ATGAGTGATGGCGCA-MGB-NFQ | | + | |
| RV | F | JVKF-v2 | CAGTGGTTGRTGCTCAAGATGGA | 131 | + | This study |
| RV | R | JVKR-v2 | TCATTRTAATCATATTGAATACCCA | | - | This study |
| RV | P | JVKP | FAM-ACAACTGCA/ZEN/GCTTCAAAAGAAGWGT-IBFQ | | - | (Jothikumar et al., 2009) |

[a] FAM, 6-carboxyfluorescein; IBFQ, Iowa Black Fluorescent quencher; MGB, minor groove binder; NFQ, nonfluorescent quencher; TAMRA, 5-carboxytetramethylrhodamine

Table S2. Basic metrics of standard curve for each virus and protozoa assay.

| **Target** | **Intercept** | **Slope** | **$R^2$** | **PCR Efficiency** | **LOQ ($10^x$ gc/reaction)** |
|---|---|---|---|---|---|
| AdV-F | 33.3 | -3.7 | 0.92 | 87% | 1.1 |
| AiV | 31.9 | -4.0 | 0.98 | 77% | 1.1 |
| SaV | 31.8 | -3.7 | 0.98 | 86% | 1.1 |
| EV | 30.9 | -3.6 | 0.99 | 89% | 1.1 |
| NoV GI | 32.1 | -3.7 | 0.99 | 85% | 2.1 |
| NoV GII | 31.7 | -4.2 | 0.99 | 74% | 2.1 |
| SARS-CoV-2 | 29.1 | -3.4 | 0.98 | 95% | 1.1 |
| IAV | 29.2 | -4.0 | 0.99 | 79% | 1.1 |
| PMMoV | 32.0 | -4.0 | 0.97 | 77% | 2.1 |
| BKPyV | 34.2 | -4.5 | 0.95 | 66% | 2.1 |
| JCPyV | 28.8 | -3.8 | 1.00 | 83% | 1.1 |
| *Carjivirus* | 30.0 | -3.2 | 0.98 | 107% | 0.1 |
| CGMMV | 31.0 | -3.8 | 1.00 | 84% | 1.1 |
| TMGMV | 29.1 | -3.9 | 0.99 | 81% | 1.1 |
| ToBRFV | 30.9 | -4.1 | 0.95 | 75% | 0.1 |
| ToMV | 32.9 | -3.2 | 0.99 | 107% | 1.1 |
| TMV | 28.9 | -3.7 | 0.99 | 86% | 1.1 |
| ToMMV | 27.5 | -3.6 | 0.98 | 91% | 0.1 |
| TSAMV | 28.6 | -4.0 | 0.99 | 79% | 1.1 |
| *Giardia* | 30.1 | -3.9 | 1.00 | 82% | 1.1 |
| MNV | 30.0 | -3.0 | 1.00 | 114% | 2.1 |

Table S3. A summary of *p*-values of Tukey's HSD *post hoc* comparison for reductions of different targets. P-values less than 0.05 were highlighted with red color.

| | **AdV-F** | **AiV** | **SaV** | ***Carjivirus*** | **PMMoV** | **BKPyV** | **JCPyV** | **CGMMV** | **TMGMV** | **ToBRFV** | **ToMV** | **TMV** | ***Giardia*** |
|---|---|---|---|---|---|---|---|---|---|---|---|---|---|
| **AdV-F** | | $1.0\times10^{0}$ | $1.0\times10^{0}$ | $8.2\times10^{-13}$ | $1.0\times10^{0}$ | $5.4\times10^{-3}$ | $2.6\times10^{-10}$ | $7.8\times10^{-1}$ | $1.0\times10^{0}$ | $1.0\times10^{0}$ | $9.5\times10^{-1}$ | $1.0\times10^{0}$ | $2.6\times10^{-1}$ |
| **AiV** | | | $1.0\times10^{0}$ | $9.1\times10^{-5}$ | $1.0\times10^{0}$ | $1.5\times10^{-1}$ | $5.8\times10^{-4}$ | $7.7\times10^{-1}$ | $1.0\times10^{0}$ | $9.9\times10^{-1}$ | $8.7\times10^{-1}$ | $1.0\times10^{0}$ | $1.0\times10^{0}$ |
| **SaV** | | | | $2.8\times10^{-4}$ | $1.0\times10^{0}$ | $2.7\times10^{-1}$ | $1.7\times10^{-3}$ | $9.0\times10^{-1}$ | $1.0\times10^{0}$ | $1.0\times10^{0}$ | $9.6\times10^{-1}$ | $1.0\times10^{0}$ | $1.0\times10^{0}$ |
| ***Carjivirus*** | | | | | $6.5\times10^{-14}$ | $2.0\times10^{-3}$ | $1.0\times10^{0}$ | $8.0\times10^{-8}$ | $1.2\times10^{-13}$ | $8.7\times10^{-12}$ | $7.7\times10^{-1}$ | $3.3\times10^{-12}$ | $7.3\times10^{-14}$ |
| **PMMoV** | | | | | | $4.4\times10^{-5}$ | $2.5\times10^{-13}$ | $1.2\times10^{-1}$ | $1.0\times10^{0}$ | $9.6\times10^{-1}$ | $3.0\times10^{-1}$ | $1.0\times10^{0}$ | $8.1\times10^{-1}$ |
| **BKPyV** | | | | | | | $4.7\times10^{-2}$ | $7.1\times10^{-1}$ | $1.3\times10^{-3}$ | $2.5\times10^{-02}$ | $4.1\times10^{-1}$ | $1.0\times10^{-3}$ | $1.2\times10^{-6}$ |
| **JCPyV** | | | | | | | | $1.0\times10^{-5}$ | $2.2\times10^{-11}$ | $2.6\times10^{-09}$ | $1.3\times10^{-06}$ | $4.1\times10^{-10}$ | $3.0\times10^{-13}$ |
| **CGMMV** | | | | | | | | | $5.4\times10^{-1}$ | $9.6\times10^{-1}$ | $1.0\times10^{0}$ | $2.7\times10^{-1}$ | $2.0\times10^{-3}$ |
| **TMGMV** | | | | | | | | | | $1.0\times10^{0}$ | $8.2\times10^{-1}$ | $1.0\times10^{0}$ | $4.0\times10^{-1}$ |
| **ToBRFV** | | | | | | | | | | | $1.0\times10^{0}$ | $9.8\times10^{-1}$ | $1.1\times10^{-1}$ |
| **ToMV** | | | | | | | | | | | | $5.0\times10^{-1}$ | $6.9\times10^{-3}$ |
| **TMV** | | | | | | | | | | | | | $9.4\times10^{-1}$ |
| ***Giardia*** | | | | | | | | | | | | | |

Table S4. Geometric mean of the influent concentration, coefficient of variation in concentration, and differences in $\log_{10}$ reduction with AdV-F and SaV. Each dataset is color-coded depending on the descending rank of geometric mean, and on the ascending rank of coefficient of variation and absolute differences in $\log_{10}$ reductions with AdV-F and SaV. The top rank is highlighted in red, while the bottom rank is highlighted in blue, with a gradient in between.

| | Abundance | Variation | Comparability | |
|---|---|---|---|---|
| | Geometric mean ($\log_{10}$ gc/L) | Coefficient of variation | Difference in reduction with AdV-F ($\log_{10}$) | Difference in reduction with SaV ($\log_{10}$) |
| **PMMoV** | 7.0 | 0.5 | −0.1 | 0.1 |
| **CGMMV** | 6.1 | 0.6 | 0.3 | 0.5 |
| **TMGMV** | 5.8 | 0.4 | −0.0 | 0.2 |
| **ToBRFV** | 6.9 | 1.4 | 0.1 | 0.3 |
| **ToMV** | 6.0 | 1.1 | 0.2 | 0.4 |
| **TMV** | 4.7 | 1.0 | −0.2 | 0.1 |
| ***Carjivirus*** | 9.1 | 0.8 | 1.1 | 1.4 |
| **BKPyV** | 6.6 | 0.4 | 0.6 | 0.8 |
| **JCPyV** | 6.5 | 0.9 | 1.0 | 1.2 |

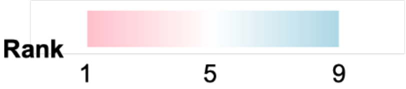
